# A protocol for high-throughput microplate-based CUT&Tag

**DOI:** 10.1101/2025.09.18.676951

**Authors:** Brittany B. Johnson, Gabriel E. Boyle, Arnold J. Federico, Jing Ma, Humza Hemani, David H. Spencer, Jay F. Sarthy, Michael P. Meers

## Abstract

Cleavage Under Targets & Tagmentation (CUT&Tag) is a versatile method for measuring genomic occupancy of chromatin-associated proteins with high sensitivity and specificity. CUT&Tag has low sequencing requirements and is therefore suitable for highly multiplexed experiments, but methods to process samples at throughput without specialized equipment are lacking. Here we present a method for simultaneous parallel processing of 96 CUT&Tag samples in a standard microplate. Plate-CUT&Tag can be carried out in a similar time frame to benchtop CUT&Tag and yields data of comparable quality. We present data from cell culture and patient leukemia samples processed with Plate-CUT&Tag to illustrate its utility in large-scale preclinical and translational studies.

## Introduction

Eukaryotes package their genomes into a protein-DNA complex known as chromatin, of which the nucleosome is the elemental subunit: a globular, hetero-octameric complex of histone proteins that stably wraps ~150 bp of DNA around its core^1,2^. Chromatin plays a role in nearly all DNA-dependent processes by acting as a default barrier to genome access, modified by myriad chromatin-associating proteins and chemical modifications of histones^3–5^. Accordingly, an accurate quantification of the enrichment of chromatin-associated proteins and modifications throughout the genome is essential for understanding how such processes are regulated. DNA sequencing-based “chromatin profiling” methods such as Chromatin Immunoprecipitation Sequencing (ChIP-seq) are the gold standard for identifying genomic sites of protein enrichment. In ChIP-seq, formaldehyde-crosslinked, solubilized chromatin is immunoprecipitated with antibodies against chromatin-associated proteins of interest, and the associated DNA is sequenced and compared with DNA from non-immunoprecipitated control chromatin^6^. More recently introduced “enzyme tethering” methods for chromatin profiling, including Cleavage Under Targets and Release Using Nuclease (CUT&RUN) and Cleavage Under Targets and Tagmentation (CUT&Tag), profile chromatin in its native context by dispensing with crosslinking and solubilization steps^7,8^. These adaptations result in high signal-to-noise and consequently lower sequencing requirements that make CUT&RUN and CUT&Tag suitable for high-throughput experimental studies, including for clinical samples and translational applications. Adaptations to CUT&RUN and CUT&Tag for operation on automated liquid handling platforms have made this a reality, with tens to hundreds of samples processed simultaneously to reduce sequencing bias^9,10^.

Nevertheless, high-throughput chromatin profiling protocols are less accessible to individual labs owing to specialized equipment requirements. To democratize high-throughput chromatin profiling, methods for highly parallel benchtop processing of samples by a single operator using standard laboratory equipment and reagents are desirable. Here we describe a simple protocol for parallel benchtop CUT&Tag in a standard 96 well microplate which we denote as “Plate-CUT&Tag”. We demonstrate Plate-CUT&Tag’s effectiveness for profiling histone post-translational modifications and non-histone chromatin-associated proteins in cell lines and patient Acute Myeloid Leukemia (AML) samples. Plate-CUT&Tag can serve as a core method for high-throughput chromatin profiling in nearly any laboratory setting.

## Results

The previously reported “one-pot” CUT&Tag-Direct protocol^11^ yields CUT&Tag libraries from cells in a single tube. We made adaptations to this protocol for use with multichannel pipettes in the wells of a 96 well microplate (Fig. 1A, Supplementary Fig. 1A). Specifically, we adjusted the volumes and concentrations of two critical steps: the SDS denaturation step after tagmentation, and the Triton neutralization step prior to PCR amplification, to improve their reliability for multichannel pipetting. After testing multiple protocol variations, we found that incorporating a larger volume of 0.1% SDS in the denaturation step combined with a higher concentration of Triton X-100 in the neutralization step resulted in more reliable yields in Plate-CUT&Tag experiments using human Chronic Myelogenous Leukemia (CML) cells (K562) and cells from two different AML patients (Supplementary Fig. 1B). In comparison with CUT&Tag-Direct, Plate-CUT&Tag requires a similar number of steps and a similar bench duration time, while enabling more samples to be processed in parallel (Fig. 1A, Supplementary Fig. 1A).

**Figure 1.**
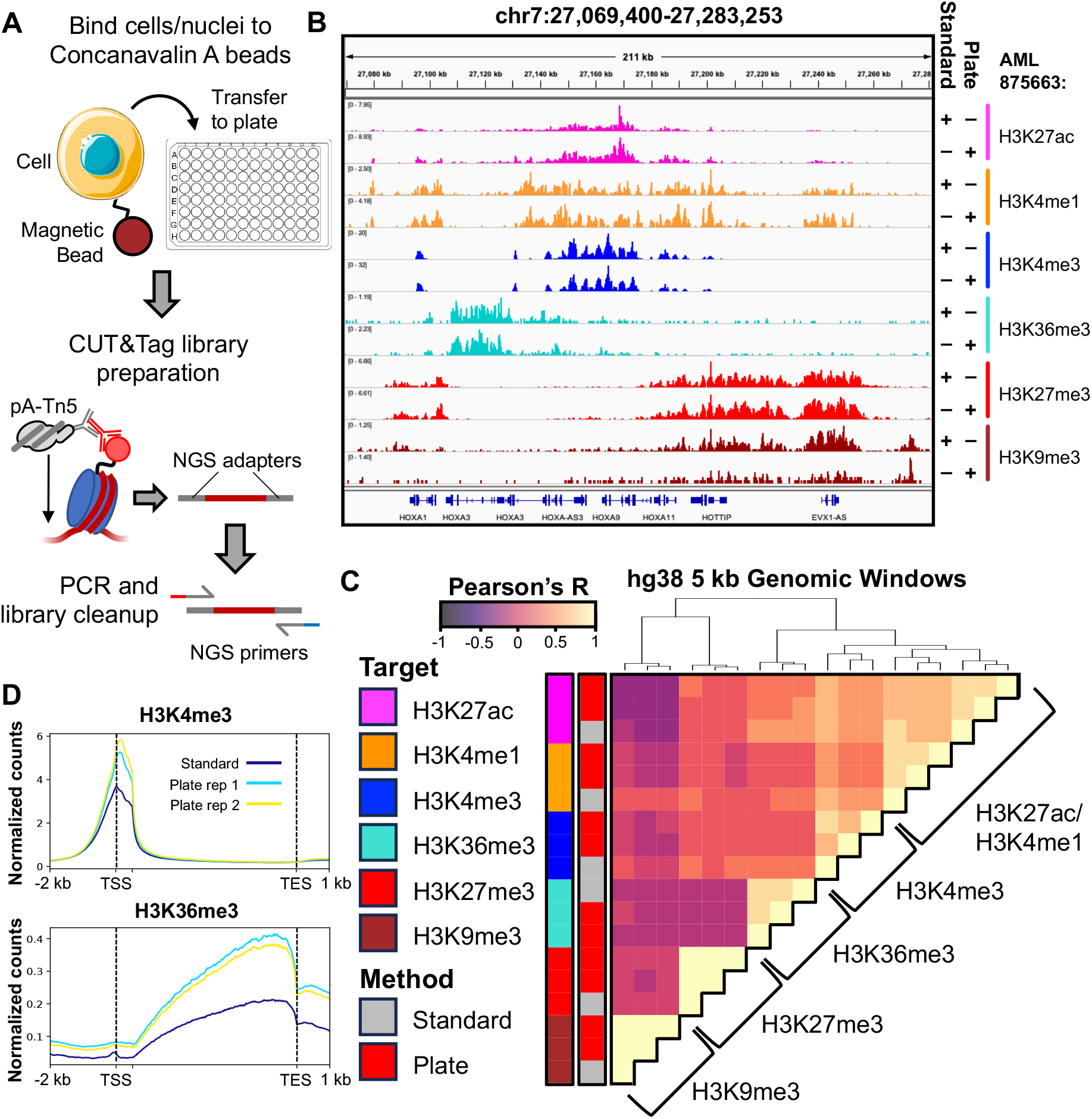
A) Schematic of Plate-CUT&Tag protocol. B) Integrative Genomics Viewer (IGV)^30^ browser screenshot depicting matched standard (CUT&Tag-Direct) or Plate-CUT&Tag datasets mapping six indicated histone post-translational modifications at the HOXA locus from a single AML patient. C) Heatmap depicting Pearson’s correlation coefficients (R) of enrichment of six indicated histone post-translational modifications in standard or Plate-CUT&Tag datasets across 5 kb windows spanning the hg38 genome. D) Average plots depicting H3K4me3 (top) and H3K36me3 (bottom) mean enrichment across hg38 gene bodies in standard (blue) or Plate-CUT&Tag (teal, yellow) experiments.

In a pilot experiment comparing the efficacy of plate-based with standard CUT&Tag for profiling samples of clinical interest, we generated one standard CUT&Tag-Direct replicate (50K nuclei) and two plate-based CUT&Tag-Direct replicates (Plate-CUT&Tag, 25K nuclei each) targeting each of six histone post-translational modifications (PTMs; H3K27ac, H3K4me1, H3K4me3, H3K36me3, H3K27me3, and H3K9me3) in AML blasts from a single patient (Supplementary Table S1). Profiles were highly similar and the majority of peaks were shared for the same target between standard and plate-based methods (Fig. 1B, Supplementary Fig. 2), although fragment duplicate rates were slightly higher in plate samples (Supplementary Table S1). To quantify concordance between standard and plate samples, for all pairs of samples we calculated a pairwise Pearson’s correlation of normalized count enrichment in 5 kb windows spanning the genome and hierarchically clustered the result. We found that there was perfect discrimination between targets in clustering and high correlation between standard and plate samples within the same target (Fig. 1C). Moreover, we observed expected genomic patterns of histone modifications in both standard and plate samples, including significant overlap of H3K27ac and H3K4me1 in regulatory regions (Fig. 1C), enrichment of H3K4me3 around promoters and TSSs, and enrichment of H3K36me3 in the 3’ ends of genes (Fig. 1D). These results indicate that Plate-CUT&Tag is equivalent to standard CUT&Tag-Direct in data quality.

Epigenomics methods are often challenging to perform on non-nucleosomal targets whose association with chromatin is transient, such as transcription factors and chromatin remodelers. To demonstrate Plate-CUT&Tag’s utility for profiling such proteins, we used Plate-CUT&Tag to profile RNA Polymerase II, chromatin remodeler subunit BRG1, insulator boundary protein CTCF, and transcription factor MYC, in K562 CML and RS4-11 Acute Lymphoblastic Leukemia (ALL) cell lines (Fig. 2A-B). All targets were profiled at high signal-to-noise with a high fraction of reads in peaks (FRiP) relative to non-targeted controls with minimal optimization necessary (Supplementary Table S2). Factors were enriched in their expected genomic patterns: for instance, RNA PolII was enriched in the gene body relative to exclusively promoter-enriched histone PTM H3K4me3, reflecting productive transcriptional elongation (Supplementary Fig. 3A). In contrast, BRG1 and Myc were highly promoter enriched, consistent with reported roles in shifting promoter-bound nucleosomes and stimulating transcription initiation, respectively^12,13^ (Supplementary Fig. 3B-C).

**Figure 2.**
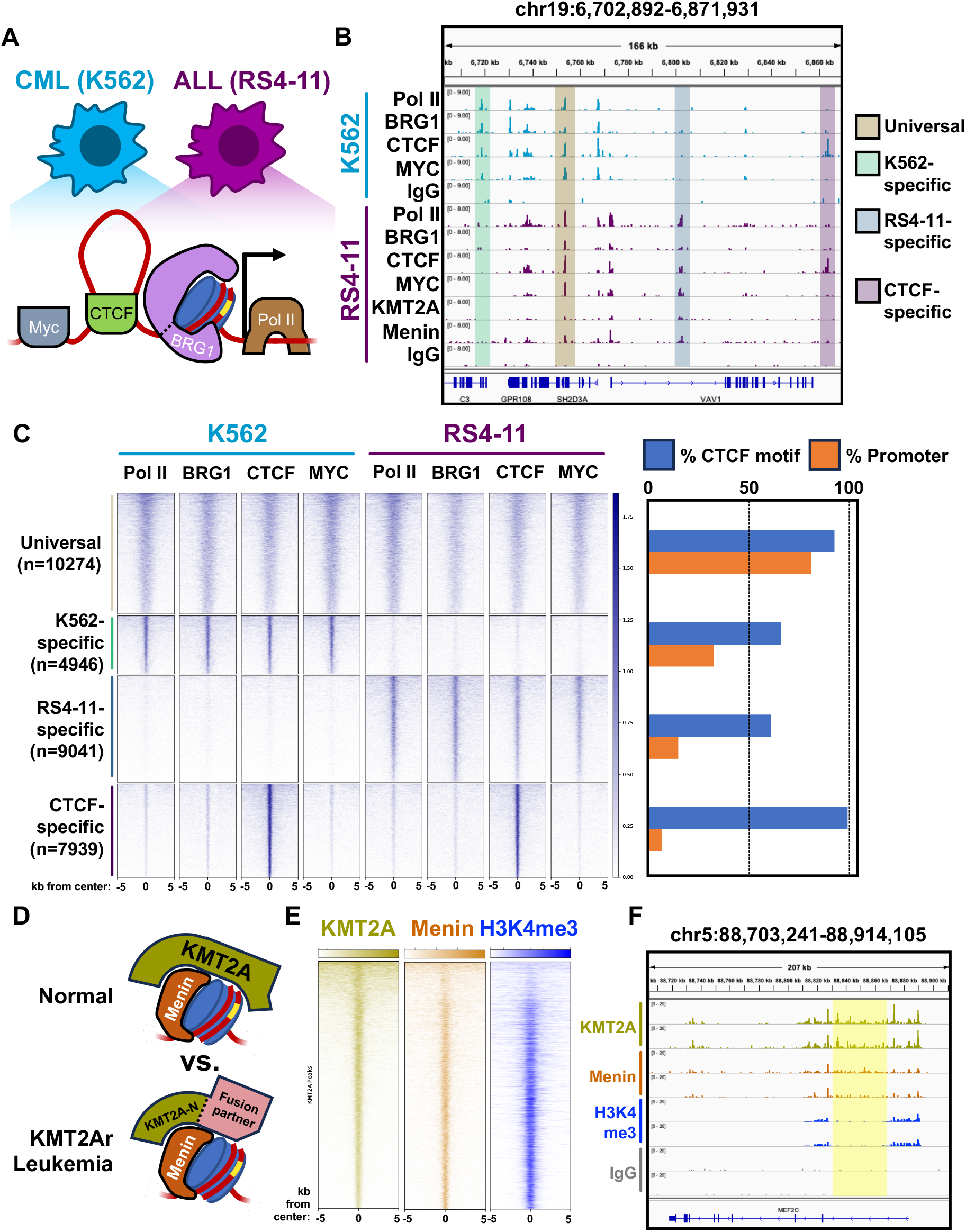
A) Schematic of profiling non-histone targets in CML and ALL cell lines. B) IGV browser screenshot depicting enrichment of indicated targets in K562 (first 5 rows) or RS4-11 (last 7 rows) cells. Regions of universal (yellow), K562-specific (green), RS4-11-specific (blue), or CTCF-specific (purple) enrichment are shaded. C) Left: Heatmap depicting enrichment of indicated targets in a 10 kb window surrounding universal (yellow), K562-specific (green), RS4-11-specific (blue), or CTCF-specific (purple) regions, as defined by K-means clustering (Methods). Right: Bar chart depicting the percentage of promoter-overlapping regions (orange) and percentage of regions containing a high-confidence CTCF motif (blue) within each cluster. D) Left: Schematic of KMT2A and Menin binding in normal and KMT2Ar leukemia. Right: Heatmap depicting KMT2A (gold), Menin (orange), and H3K4me3 (blue) enrichment in a 10 kb window around KMT2A peaks in RS4-11 cells. F) IGV browser screenshot depicting enrichment of indicated targets at the MEF2C locus in RS4-11 cells. Region of KMT2A and Menin enrichment in the absence of H3K4me3 enrichment is shaded yellow.

We identified enriched peaks for each non-histone factor and calculated pairwise Pearson correlation coefficients based on their enrichment in a merged peak set (n=32200). We found that they clustered primarily by their cell type of origin, consistent with cell type-specific regulatory patterns (Supplementary Fig. 3D). This was recapitulated on the locus level, where K-means clustering (Methods) partitioned most individual peaks into clusters that corresponded to K562- or RS4-11-specific sites, in addition to a “universal” cluster with shared enrichment across cell types (Fig. 2C). A fourth cluster was enriched specifically for CTCF from both cell types, representing cell type-agnostic insulators characterized by a high incidence of CTCF motifs and residence in distal regions outside of promoters (Fig. 2C). We went on to conduct enriched motif analysis from sequences derived from called peaks and found that the preferred CTCF binding motif is the most highly enriched in CTCF peaks, indicating Plate-CUT&Tag accurately captures sequence-specific CTCF binding (Supplementary Fig. 3E). Curiously, the preferred E-Box motif of Myc (CANNTG) was not detected as a top motif in Myc-enriched peaks in either cell line, though this is potentially consistent with a reported role of nonspecific promoter DNA binding in coordinating Myc’s regulatory function^14^ (Supplementary Fig. 2D). BRG1 is not reported to engage in sequence-specific binding, but nevertheless we uncovered enriched motifs for FOS/Jun-family factors (SP1, SP2, SP4), NRF1, and RUNX1 that likely reflect core regulatory factors with which BRG1 binds in these cell types (Supplementary Fig. 3E).

To demonstrate that Plate-CUT&Tag can profile complex regulatory interactions between protein binding partners, we turned to the RS4-11 ALL cell line. RS4-11 originated from an ALL patient bearing a t(4;11)(q21;q23) translocation that produces a breakpoint fusion between the N-terminus of the *MLL1* gene encoding the histone methyltransferase KMT2A, and the C-terminus of the *AFF1* gene encoding the transcriptional elongation factor AF4, resulting in a KMT2A-AF4 “oncofusion” protein. t(4;11) translocations between KMT2A and a variety of different fusion partners define the KMT2A-rearranged (KMT2Ar) subtype of leukemia, patients of which are often refractory to frontline treatment and exhibit high rates of relapse^15^. KMT2A-AF4 localizes to chromatin and drives aberrant enhancer activation and gene expression outcomes in ALL^10,16^. To demonstrate the utility of Plate-CUT&Tag for profiling therapeutically important proteins, we used RS4-11 cells to profile KMT2A and its binding partner, the scaffolding protein Menin, which is a target for small-molecule inhibitors used to treat KMT2Ar leukemias^17^ (Fig. 2D). In RS4-11 cells, KMT2A binding is highly concordant with that of Menin, and is enriched at active promoters associated with H3K4me3 as previously reported^10^ (Fig. 2E, Supplementary Fig. 3F). Interestingly, sites most highly enriched for KMT2A often exhibit reduced H3K4me3 (Fig. 2E, top of plot), which may correspond to previously reported regions in which KMT2A oncofusion ingresses into the gene bodies of key hematopoietic regulatory genes^10^ (Fig 2F). In all, these data demonstrate that Plate-CUT&Tag can profile molecularly and therapeutically important regulatory factors that interact indirectly with chromatin.

To demonstrate Plate-CUT&Tag’s ability to carry out large-scale clinical studies, we profiled the aforementioned six histone PTMs across ten patient AMLs representing distinct mutational backgrounds in biological duplicates, for a total of 120 Plate-CUT&Tag experiments across two 96 well plates (Fig. 3A). The experiments in their entirety were accomplished in roughly four working days by a single technician. Principal component analysis (PCA) of the resulting enrichment profiles showed that experiments clustered dominantly by PTM rather than by patient identity within PCA space, consistent with broadly shared, PTM-specific genomic enrichment in samples from blood/bone marrow with high regulatory similarity (Fig. 3B, Supplementary Fig. 4A). Concordantly, principal component 1 (27.29% of variance) separated PTMs by “active” (H3K27ac, H3K4me1, H3K4me3, H3K36me3) or “repressive” (H3K27me3, H3K9me3) character whose enrichment tend to occur in mutually exclusive genomic regions (Fig. 3B). Nevertheless, variability within PTMs was indicative of patient-specific signatures. For instance, in one patient, H3K9me3 profiles clustered in the H3K27me3 PCA domain (Fig. 3B, black arrows); genome browser and shared enrichment analysis confirmed that H3K27me3 and H3K9me3 profiles were highly co-enriched in this patient alone (Supplementary Fig. 4B-C).

**Figure 3.**
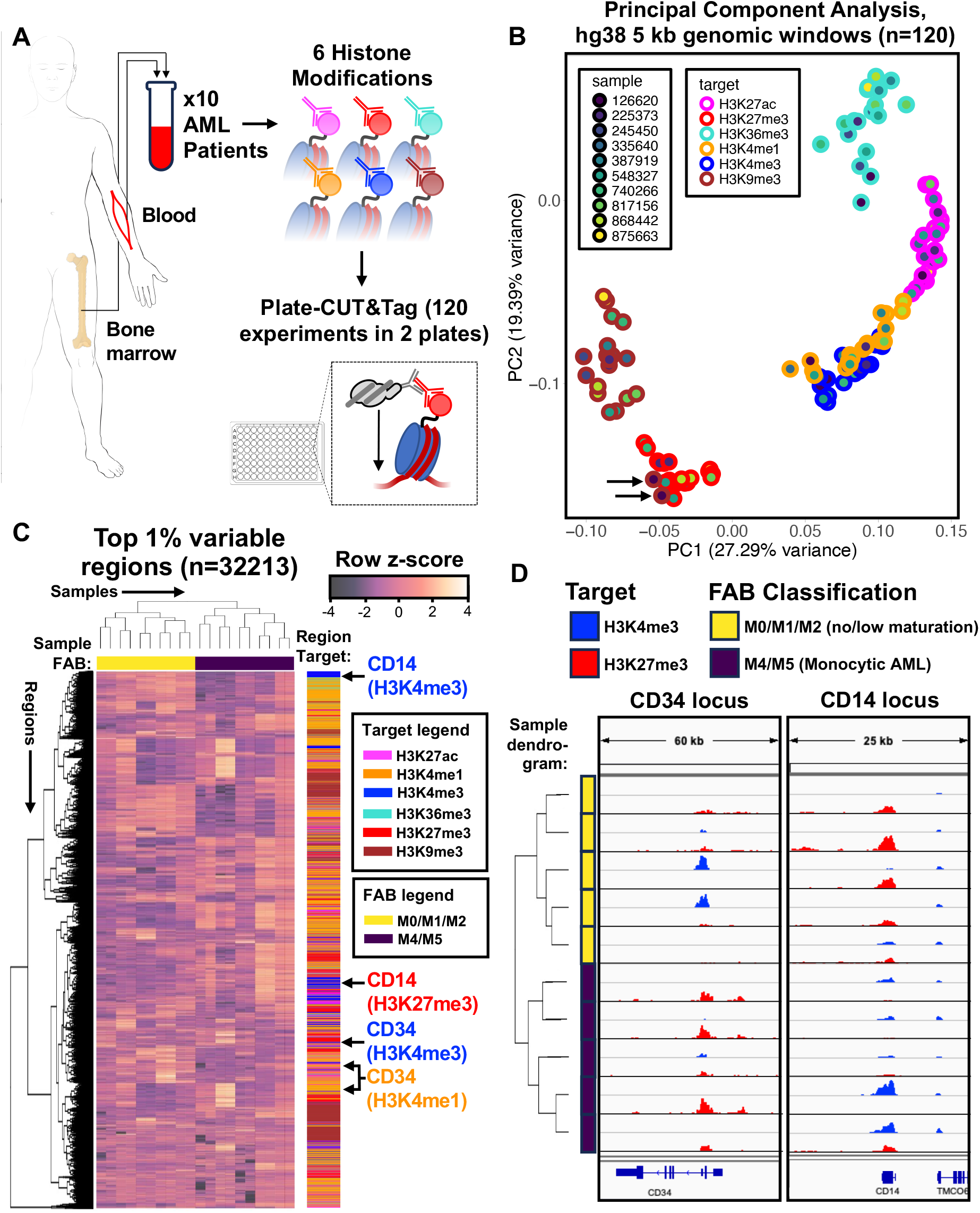
A) Schematic of pilot experiment profiling six histone post-translational modifications in ten AML patient samples via Plate-CUT&Tag. B) Principal Component Analysis (PCA) of enrichment in 5 kb windows spanning the hg38 genome from 120 Plate-CUT&Tag datasets representing all combinations of target, patient of origin, and replicate. Black arrows denote patient in which H3K9me3 datasets cluster closely with H3K27me3. C) Heatmap displaying Plate-CUT&Tag enrichment for all targets in all patients in the top 1% most variable genomic regions as denoted by standard deviation. Samples are colored by French-American-British (FAB) Classification on the X-axis, and regions are colored by target of origin on the Y-axis. D) IGV genome browser screenshot depicting enrichment of H3K4me3 (blue) and H3K27me3 (red) across all patients at the CD34 (left) and CD14 (right) loci. Samples are colored by French-American-British (FAB) Classification on the Y-axis.

To illustrate our ability to glean patient-specific, locus-level insights from high dimensional Plate-CUT&Tag data, we co-embedded normalized count matrices for each PTM in the same samples so that PTM-specific features could be compared to each other (Methods). We then calculated feature-specific standard deviations across patients to identify the features contributing to differences between patients, and selected features in the top 1% of standard deviation for further analysis (n=32213) (Fig. 3C, Supplementary Fig. 4D). Notably, among the most variable features were H3K4me1/3 and H3K27me3 at the CD34 and CD14 loci, expression markers of hematopoietic stem cells (HSCs) and monocytes, respectively^18,19^ (Fig. 3C-D). Upon closer inspection, we found that within individual patients, CD34 and CD14 were enriched for H3K4me3 and H3K27me3 in a mutually exclusive manner, perhaps indicative of clonal leukemia that is largely either HSC-like (CD34-expressing) or monocyte-like (CD14-expressing) (Fig. 3D). Indeed, samples exhibiting CD34 H3K4me3 enrichment and/or CD14 H3K27me3 enrichment were classified as exhibiting no or limited maturation by the FAB Classification system (M0/M1/M2), consistent with CD34-expressing AML without maturation or with minimal differentiation^20,21^ (Fig. 3C-D). Similarly, samples with CD14 H3K4me3 and/or CD34 H3K27me3 were all classified as monocytic AML (M4/M5) (Fig. 3C-D). These results show that Plate-CUT&Tag can be used to identify clinically relevant molecular phenotypes on the locus level across many patients in the same experiment.

## Discussion

We have introduced here a reliable method for conducting highly parallel CUT&Tag in 96-well microplates. Parallelized protocols for genomics techniques are highly desirable to minimize technical error across samples commonly introduced by PCR or sequencing. However, such approaches commonly use robotics or other automated systems that require high-cost equipment/reagents and specialized expertise to operate. In contrast, Plate-CUT&Tag can accomplish 96 CUT&Tag experiments by a human operator using standard lab reagents and equipment in a similar time frame to standard CUT&Tag-Direct. Thus, Plate-CUT&Tag can democratize CUT&Tag for highly parallel experiments for individual research labs that cannot afford or otherwise use specialized automated platforms.

We conducted a pilot experiment using Plate-CUT&Tag to profile histone post-translational modifications in 10 AML patient samples that are challenging to profile by conventional means. When deployed at scale in models of disease via Plate-CUT&Tag, chromatin profiling is capable of uncovering novel molecular insights unattainable via other methods that might bear on future therapeutic interventions. For instance, we found that H3K4me1 features were dominant contributors to patient-specific stratification of AML samples (Supplementary Fig. 4D). H3K4me1 is notably a marker of poised or minimally active enhancers^22–24^, suggesting that regulatory characteristics that aren’t captured by the genes expressed via fully active enhancers/promoters may be important classifiers of disease. We also show that difficult-to-profile targets, including transcription factors and chromatin modifiers, are well within reach using Plate-CUT&Tag. This characteristic is of particular importance for diseases such as KMT2Ar leukemia whose etiology stems largely from the aberrant function of DNA binding proteins. In all, Plate-CUT&Tag present a promising, highly accessible method for clinical-scale chromatin profiling.

### Limitations

Because the hands-on bench time involved in Plate-CUT&Tag is comparable to CUT&Tag-Direct, it does not achieve the same savings in hands-on time as protocols that are fully automated by liquid handling systems. Plate-CUT&Tag is therefore likely to be most useful for researchers interested in processing many samples in parallel, but who lack the resources or infrastructure to establish an automated liquid handling system in-house. We also did not explicitly test the efficacy of Plate-CUT&Tag on permeabilized whole cells, which might be a particularly useful application with limited clinical samples. Based on our success using CUT&Tag-Direct on whole cell samples in-house (data not shown), we expect that Plate-CUT&Tag would be suitable for such samples. We have also found in our pilot studies that Plate-CUT&Tag samples have slightly higher duplicate rates than matched samples processed by benchtop CUT&Tag-Direct (Supplementary Table S1). This may be due to insufficiently optimized SDS release and Triton X-100 neutralization steps, which are essential for denaturing Tn5 to release tagmented fragments and neutralizing SDS before PCR, respectively. Inability to carry out these steps effectively can lead to lower library yield, which might result in more PCR duplicates. The latest protocols for CUT&Tag-Direct utilize higher percentages of both SDS and Triton X-100 to ensure denaturation of highly crosslinked samples (https://dx.doi.org/10.17504/protocols.io.x54v9mkmzg3e/v5), and it is possible that such adaptations might improve Plate-CUT&Tag yields.

## Methods

### Cell culture and nuclei preparation

Human K562 chronic myelogenous leukemia cells (American Type Culture Collection (ATCC)) and RS4-11 Acute Lymphoblastic Leukemia Cells (ATCC) were authenticated for STR, sterility, human pathogenic virus testing, mycoplasma contamination and viability at thaw. K562 and RS4-11 cells were cultured in liquid suspension; the former in IMDM (ATCC) with 10% FBS added (Seradigm), and the latter in RPMI (ATCC) with 10% FBS added. K562 and RS4-11 cells were harvested by centrifugation for 3 minutes at 1,000*g* and then resuspended in 1× PBS. Lightly cross-linked nuclei were prepared from cells as described in steps 2–14 of the Bench Top CUT&Tag protocol on protocols.io (https://doi.org/10.17504/protocols.io.bcuhiwt6). In brief, cells were pelleted for 3 minutes at 600*g*, resuspended in hypotonic NE1 buffer (20 mM HEPES-KOH pH 7.9, 10 mM KCl, 0.5 mM spermidine, 10% Triton X-100 and 20% glycerol) and incubated on ice for 10 minutes. The mixture was pelleted for 4 minutes at 1,300g, resuspended in 1× PBS and fixed with 0.1% formaldehyde for 2 minutes before quenching with 60 mM glycine. Nuclei were counted using the ViCell Automated Cell Counter (Beckman Coulter) and frozen at −80 °C in 10% DMSO at a concentration of 1M nuclei/mL for future use.

### Acute Myeloid Leukemia patient sample preparation

Blood and bone marrow samples were collected at presentation from adult patients with de novo AML on an Institutional Review Board–approved banking protocol (#201011766). Nuclei were extracted from samples according to the nuclei preparation protocol listed above with the following modifications: NE1 hypotonic lysis buffer incubation time was optimized for each individual patient sample based on percentage of trypan blue-positive nuclei yield from input cells, with all samples incubated for between 45 seconds-2 minutes. All patient samples were preserved as native nuclei without formaldehyde crosslinking at a maximum concentration of 1M nuclei/mL. Specific samples for which fewer than 1M nuclei were recovered were preserved at a concentration lower than 1M nuclei/mL.

### Antibodies

Antibodies used for CUT&Tag in this study were as follows: rabbit anti-H3K27me3 (Cell Signaling Technologies, CST9733S, lot 19, 1:100 dilution), rabbit anti-H3K27ac (Abcam, ab4729, lot 1059037-2, 1:100 dilution), rabbit anti-H3K4me3 (Active Motif, 39915, lot 29620008-11, 1:100 dilution; or Active Motif, 39159, 22341155-11, 1:50 dilution), mouse anti-H3K36me3 (Active Motif, 61021, lot 23819012, 1:100 dilution), rabbit anti-H3K9me3 (Abcam, ab8898, lot GR3302452-1, 1:100 dilution), rabbit anti-H3K4me1 (EpiCypher, 13-0040, lot 2134006-02, 1:100 dilution), rabbit anti-BRG1 (Abcam, ab110641, lot 1039568-24, 1:25 dilution) rabbit anti-CTCF (Abcam, ab128873, lot gr3284310-26, 1:25 dilution), rabbit anti-c-Myc/N-Myc (Cell Signaling Technologies, 13987S, lot 6, 1:25 dilution), rabbit anti-MLL1 Amino-terminal antigen (Cell Signaling Techhnologies, 14689, lot 1, 1:25 dilution), rabbit anti-Menin (Fortis Life Sciences, A300-105A, lot 12, 1:25 dilution), rabbit anti-RPB3 (Thermo Fisher, A303-771A, lot 2, 1:25 dilution), rabbit isotype control (Abcam, ab172730, lot 1050775-22, 1:50 dilution), guinea pig anti-rabbit (Antibodies Online, ABIN101961, 1:100 dilution) and rabbit anti-mouse (Abcam, ab46450, 1:100 dilution).

### CUT&Tag

CUT&Tag-Direct was carried out as previously described^11^ (https://doi.org/10.17504/protocols.io.bcuhiwt6). In brief, nuclei were thawed and bound to washed paramagnetic concanavalin A (ConA) beads (Bangs Laboratories) at a concentration of 5 µL washed beads per 50K nuclei, and then incubated with primary antibody at 4 °C overnight in Wash Buffer (10 mM HEPES pH 7.5, 150 mM NaCl, 0.5 mM spermidine and Roche Complete Protease Inhibitor Cocktail) with 2 mM EDTA. Bound nuclei were washed and incubated with 1:100 secondary antibody for 1 hour at room temperature and then washed 2x and incubated in Wash-300 Buffer (Wash Buffer with 300 mM NaCl) with 1:50 5 µM loaded pA–Tn5 for 1 hour at room temperature. Nuclei were washed 2x and tagmented in Wash-300 Buffer with 10 mM MgCl2 for 1 hour at 37 °C and then resuspended sequentially in 50 µl of 10 mM TAPS and 5 µl of 10 mM TAPS with 0.1% SDS and incubated for 1 hour at 58 °C. The resulting suspension was mixed well with 16 µl of 0.9375% Triton X-100, and then 2 µL each of F and R primers (sequences listed in Supplementary Table S3) and 25 µL 2× NEBNext Master Mix (New England Biolabs) were added for direct amplification with the following conditions: (1) 58 °C for 5 minutes, (2) 72 °C for 5 minutes, (3) 98 °C for 30 seconds, (4) 98 °C 15 seconds, (5) 60 °C for 15 seconds, (6) repeat steps 4–5 13 times, (7) 72 °C for 1 minute and (8) hold at 8 °C. DNA from amplified product was purified using 1.1× ratio of HighPrep PCR Cleanup System (MagBio) and resuspended in 25 µl of 10 mM Tris-HCl with 1 mM EDTA, and concentration was quantified using the Qubit (Thermo Fisher Scientific) and TapeStation (Agilent) systems.

### Plate-CUT&Tag

A detailed Plate-CUT&Tag protocol can be found here: dx.doi.org/10.17504/protocols.io.n2bvjed5wgk5/v1. Briefly, CUT&Tag-Direct experiments were carried out in individual wells of twin.tec lo-bind 96-well PCR microplates (Eppendorf) using multi-channel pipettes and reservoirs, with the following modifications: After tagmentation and washing with 10 mM TAPS, beads were resuspended individually well-by-well in 16 µL 10 mM TAPS with 0.1% SDS using a single-channel pipette with vigorous pipette mixing. The plate was sealed and vortexed briefly for 10 seconds at 1000 RPM on a Thermomixer (Eppendorf). The sealed plate was spun briefly in a swinging bucket rotor, then denatured on a 96 well deep block thermal cycler (BioRad) for 1 hour at 58^°^C. After denaturation, samples were neutralized with 5 µL 10% Triton X-100, and the plate was re-sealed and vortexed briefly for 10 seconds at 1000 RPM on the Thermomixer. Primers and NEBNext 2x Master Mix were then added, mixed, and the plate amplified according to the CUT&Tag-Direct thermal cycler program on a 96 well block. Libraries were purified and eluted as described above.

### Sequencing and data pre-processing

Libraries were sequenced on an Illumina NovaSeq X instrument with paired-end 150 × 150 reads. Raw FASTQ files were adapter trimmed using fastp^25^ and subsequently were aligned to the UCSC hg38 genome build using Bowtie2^26^, version 2.4.2, with parameters –end-to-end–very-sensitive–no-mixed–no-discordant -q–phred33 -I 10 -X 700. Mapped reads were filtered using samtools^27^ view with options -b -F 12, converted to paired-end BED files containing coordinates for the termini of each read pair, and then converted to bedGraph files using BEDTools^28^ genomecov with parameter –bg. Bedgraphs were converted to BigWig files using the UCSC bedgraphToBigwig^29^ function for visualization in the Integrative Genomics Viewer Genome Browser^30^. Raw read counts, alignment rates, and duplicate rates for all sequencing datasets presented in this study are listed in Supplementary Table S1.

### Data Analysis

#### Data and Code availability

All raw and processed data from this manuscript is deposited in the Gene Expression Omnibus (GEO) repository under Accession Number GSE308505. Code sufficient to reproduce the major analyses in this paper can be found here: https://github.com/MeersLabWashU/PlateCUTnTag.

#### Count matrices and Principal Component Analyses

For correlation/concordance analyses for histone modifications, we generated a BED file that split the hg38 genome into 5 kb windows using bedtools window and generated six modification-specific count matrices across all samples by intersecting the 5 kb windows BED file with sample paired-end fragment BED files using bedtools intersect -c. The bottom 40% of window entries by sum of counts in the row were removed and the resulting counts matrix was log10-transformed and then z-score-transformed using the *scale* utility in base R. For multi-modifications analyses from Fig. 3, matrices were generated as follows: all rows in all matrices for which row count sum was in the bottom 40% of any individual matrix was removed, and chrX, chrY, and chrM entries were removed, resulting in six matrices each with an equal number of rows and columns. For the comparison across modifications in Fig. 3B and Supplementary Fig. 4C, the six modification-specific raw count matrices were joined by column and then jointly log and z-score transformed. For the comparison across samples in Fig. 3C, each modification-specific count matrix was independently transformed, and then the resulting matrices were joined by row. Principal component analysis was performed directly on count matrices prepared as described above using the *prcomp* utility in base R.

#### Heatmaps

Correlation and variable feature heatmaps were generated using *heatmap*.*2* from the *gplots* library. For correlation heatmaps in Fig. 1C, Supplementary Fig. 3D, and Supplementary Fig. 4C, the *cor* utility in R was used to generate a correlation matrix that served as a direct input to the heatmap. For the feature heatmap in Fig. 3C, the top 1% of rows with the highest standard deviation as measured by the *sd* utility in R were plotted. Heatmaps scale was displayed using color palettes from the Viridis library.

#### Average and enrichment plots

Average and enrichment plots were generated using deeptools^31^ with functions computeMatrix, plotProfile (average plots) and plotHeatmap (enrichment plots). For average and enrichment plots scaled to gene length as in Fig. 1D, Supplementary Fig. 3A-C, and Supplementary Fig. 4A, we used computeMatrix scale-regions with the following options: --missingDataAsZero --regionBodyLength 5000 -b 2000 -a 1000 -a 1000 --unscaled5prime 500. For average and enrichment plots centered on peaks as in Figs. 2C and 2E, we used computeMatrix reference-point with the following options: --referencePoint center --missingDataAsZero -b 5000 -a 5000.

#### Peak calling

Peak calling was performed using SEACR^32^ v1.4 using a shared IgG control dataset with the following options: -n norm -m relaxed -e 0.1. For non-histone datasets, we used Irreproducible Discovery Rate (IDR)^33^ calculations to obtain a consensus set of high-confidence peaks shared between replicates for each sample-target combination. We then used bedops^34^ -m to obtain a union set of peaks from all IDR-selected peak sets for all non-histone targets shared between samples (RNA PolII, BRG1, CTCF, and c-Myc).

#### K-means clustering

K-means clustering of non-histone peaks was performed using the *kmeans* utility in base R. To determine the optimal number of initial clusters to select, we used the “knee plot” method to identify the bend in the curve plotted by number of clusters vs. within-cluster sum of squares, which occurred at 4 clusters.

#### Motif analysis

For all motif analyses, analyzed peak sets were converted to FASTA files using bedtools getfasta. For *de novo* motif discovery in Supplementary Fig. 3E, peak FASTA files were analyzed via MEME-ChIP^35^ using default options. For CTCF motif analysis in Fig. 2C, relevant peak FASTA files were scanned for the CTCF position weight matrix MA0139.1 from JASPAR using FIMO^36^ with standard options, and the percentage of peaks containing at least one motif meeting the threshold was calculated.

## Author Contributions

B.B.J.: Conceptualization, Methodology, Investigation, Validation. G.E.B: Methodology, Investigation, Validation. A.J.F.: Software. J.M.: Formal analysis, Data Curation. H.H.: Formal analysis. D.H.S.: Resources, Writing – Review & Editing. J.F.S.: Resources, Writing – Review & Editing, Supervision, Project administration, Funding acquisition. M.P.M.: Conceptualization, Methodology, Formal analysis, Resources, Data Curation, Writing – Original Draft, Writing – Review & Editing, Visualization, Supervision, Project administration, Funding acquisition.

## Acknowledgements

Thanks to D. Spencer and J. Sarthy for critical reading of the manuscript. This work was supported by a National Institutes of Health (NIH) grant to M.P.M. (R00 GM140251) and a Cancer Research Foundation Young Investigator Award grant to M.P.M.

## Supplementary Figure Legends

**Supplementary Figure 1.**
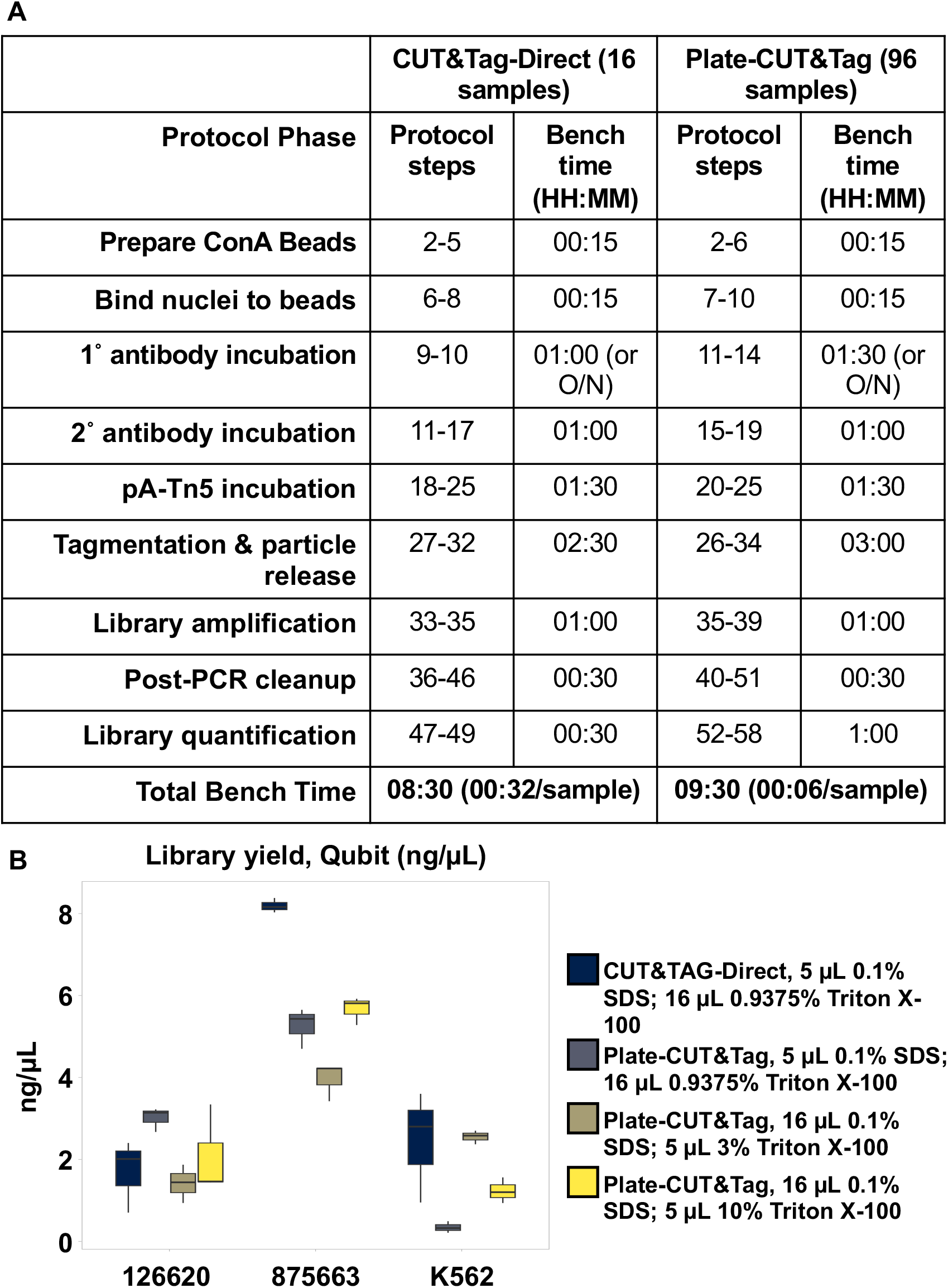
A) Table comparing the bench time spent to accomplish CUT&Tag-Direct for 16 samples vs. Plate-CUT&Tag for 96 samples. B) Boxplot denoting the DNA yield (ng/µL) obtained from libraries generated via H3K27me3 CUT&Tag in two AML patient samples (left, center) or in K562 cells (right). Colors denote the protocol used: conventional CUT&Tag-Direct (Blue), Plate-CUT&Tag using conventional CUT&Tag-Direct volumes for SDS and Triton (Grey), or Plate-CUT&Tag using two adaptations of the CUT&Tag-Direct protocol volumes for SDS and Triton X100 (Tan and Yellow). Conditions denoted in yellow were used for subsequent Plate-CUT&Tag experiments.

**Supplementary Figure 2.**
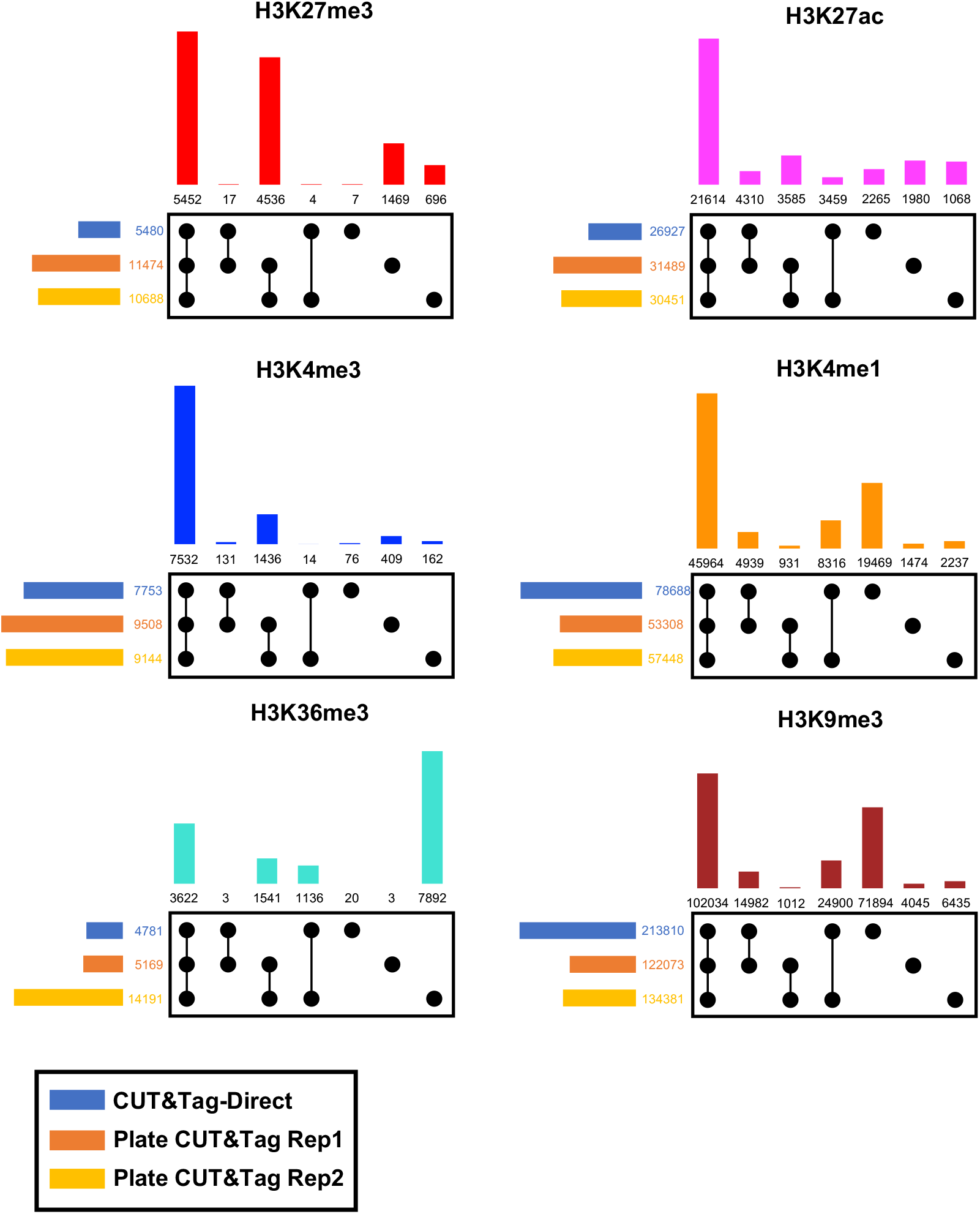
Upset plots denoting peak overlaps (top barplot) and total peaks in sample (left-hand barplot) between one CUT&Tag-Direct replicate and two Plate-CUT&Tag replicates for six histone modifications profiled in AML patient sample 875663. All overlap bar plots are ordered on the x-axis as follows from left to right: three-way intersection, Direct-Plate rep1 intersection, Plate rep1-Plate rep2 intersection, Direct-Plate rep2 intersection, Direct-specific, Plate rep1-specific, Plate rep2-specific. All peak barplots are ordered on the y-axis as follows from top to bottom: CUT&Tag-Direct, Plate-CUT&Tag rep1, Plate-CUT&Tag rep2.

**Supplementary Figure 3.**
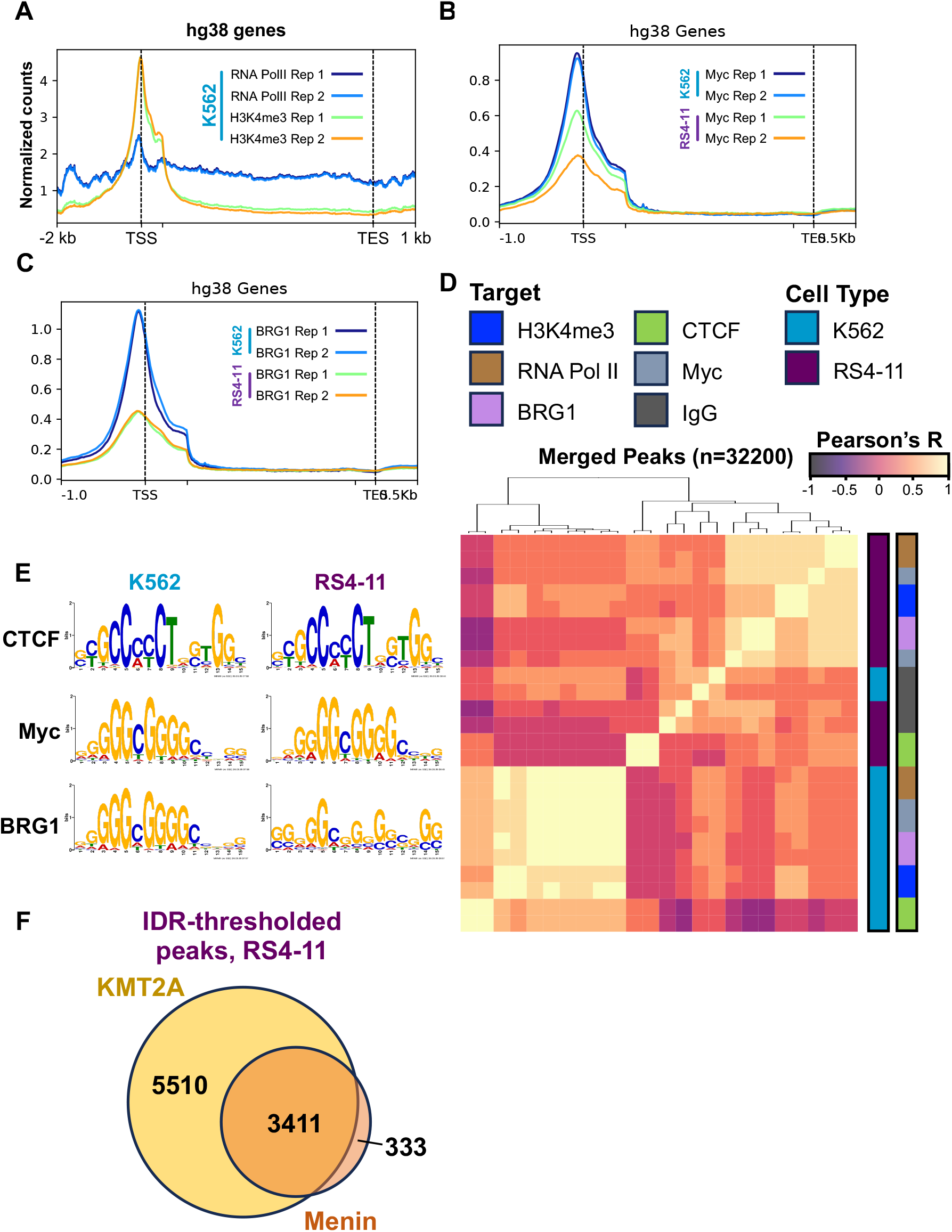
A) Average plot depicting RNA PolII (Blue, Light Blue) and H3K4me3 (Green, Orange) Plate-CUT&Tag mean enrichment across hg38 gene bodies. B) Average plot depicting Myc Plate-CUT&Tag mean enrichment across hg38 gene bodies in K562 (Blue, Light Blue) and RS4-11 (Green, Orange) cells. C) Average plot depicting BRG1 Plate-CUT&Tag mean enrichment across hg38 gene bodies in K562 (Blue, Light Blue) and RS4-11 (Green, Orange) cells. D) Heatmap depicting Pearson’s correlation coefficients (R) of enrichment of H3K4me3 (Blue) and non-histone targets RNA PolII (Brown), BRG1 (Lavender), CTCF (Green), Myc (Slate), and IgG control (Dark Grey) Plate-CUT&Tag datasets in a merged set of non-histone target peaks in K562 (Blue) and RS4-11 (Purple) cells. E) Top enriched motifs detected in CTCF (Top), Myc (Middle), and BRG1 (Bottom) IDR-filtered peaks in K562 (left) and RS4-11 (right) cells. F) Venn diagram depicting shared overlap between IDR-filtered peaks for KMT2A (Yellow) and Menin (Orange) in RS4-11 cells.

**Supplementary Figure 4.**
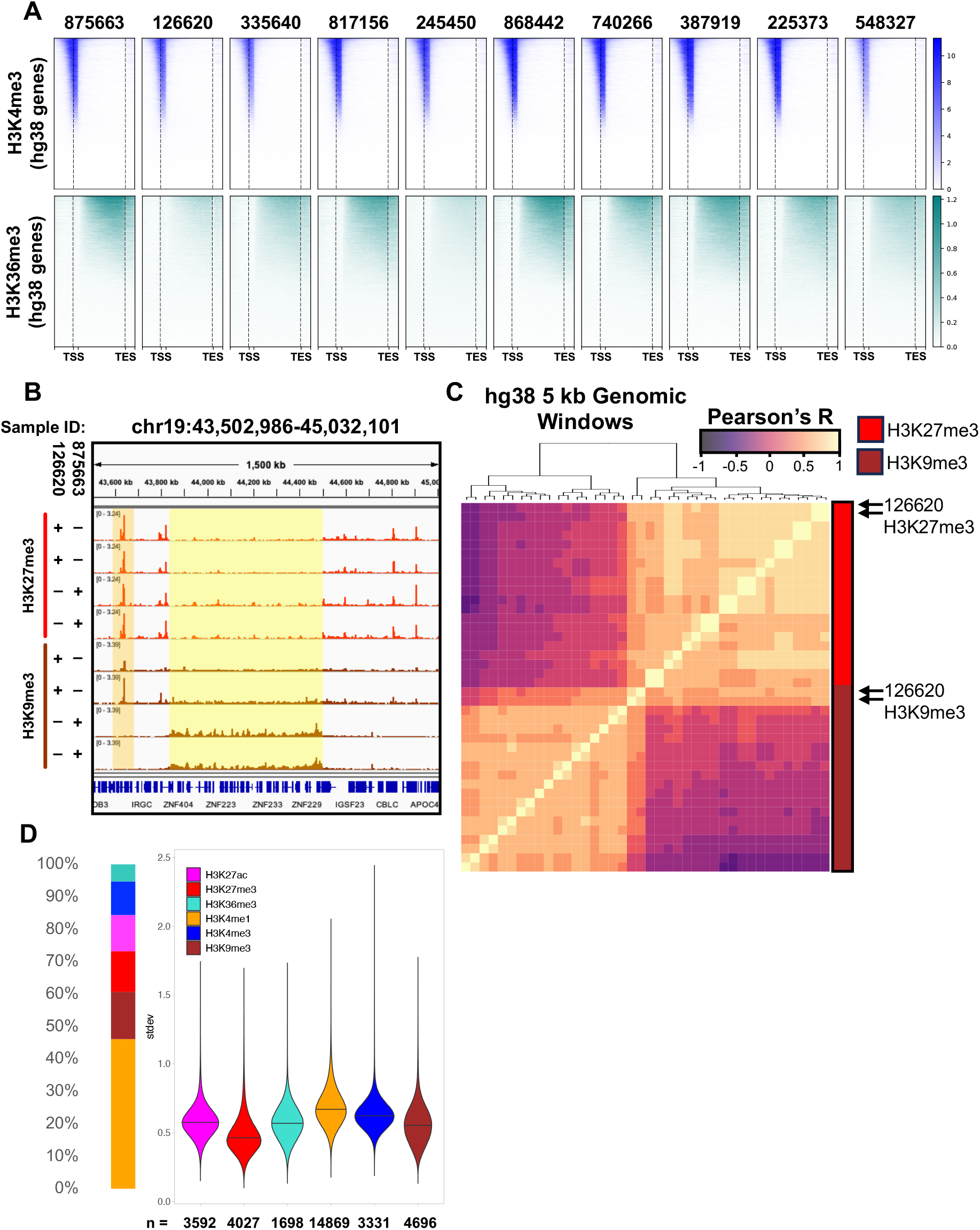
A) Heatmaps depicting H3K4me3 (Blue, Top) and H3K36me3 (Teal, Bottom) enrichment across all hg38 gene bodies for each of the 10 AML patient samples profiled in this study (IDs listed at top). Genes are ordered by modification-specific mean enrichment across all samples. B) IGV browser screenshot depicting enrichment of H3K27me3 (Top 4 rows, Red) and H3K9me3 (Bottom 4 rows, Brown) at the indicated 1.5 Mb genomic region in AML-126620 or AML-875663. C) Heatmap depicting Pearson’s correlation coefficients (R) of enrichment of H3K27me3 (Red) or H3K9me3 (Brown) across 5 kb windows spanning the hg38 genome in 10 AML patient samples. H3K27me3 and H3K9nme3 in AML-126620 are indicated by arrows. D) Left: Stacked bar plot indicating the relative proportions of features from six indicated histone modifications that are included in the top 1% of most variable features across 10 AML patient samples. Right: Violin plot depicting the distribution of standard deviations of enrichment across 10 AML patient samples (Y-axis) for features from the top 1% of variable regions, grouped by the indicated histone modifications.

## References

1 Luger, K., Mader, A. W., Richmond, R. K., Sargent, D. F. & Richmond, T. J. Crystal structure of the nucleosome core particle at 2.8 A resolution. Nature 389, 251–260, doi:10.1038/38444 (1997).

2 McGinty, R. K. & Tan, S. Nucleosome structure and function. Chem Rev 115, 2255–2273, doi:10.1021/cr500373h (2015).

3 Spitz, F. & Furlong, E. E. Transcription factors: from enhancer binding to developmental control. Nat Rev Genet 13, 613–626, doi:10.1038/nrg3207 (2012).

4 Bannister, A. J. & Kouzarides, T. Regulation of chromatin by histone modifications. Cell Res 21, 381–395, doi:10.1038/cr.2011.22 (2011).

5 Klemm, S. L., Shipony, Z. & Greenleaf, W. J. Chromatin accessibility and the regulatory epigenome. Nat Rev Genet 20, 207–220, doi:10.1038/s41576-018-0089-8 (2019).

6 Park, P. J. ChIP-seq: advantages and challenges of a maturing technology. Nat Rev Genet 10, 669–680, doi:10.1038/nrg2641 (2009).

7 Skene, P. J. & Henikoff, S. An efficient targeted nuclease strategy for highresolution mapping of DNA binding sites. Elife 6, doi:10.7554/eLife.21856 (2017).

8 Kaya-Okur, H. S. et al. CUT&Tag for efficient epigenomic profiling of small samples and single cells. Nat Commun 10, 1930, doi:10.1038/s41467-019-09982-5 (2019).

9 Janssens, D. H. et al. Automated in situ chromatin profiling efficiently resolves cell types and gene regulatory programs. Epigenetics Chromatin 11, 74, doi:10.1186/s13072-018-0243-8 (2018).

10 Janssens, D. H. et al. Automated CUT&Tag profiling of chromatin heterogeneity in mixed-lineage leukemia. Nat Genet 53, 1586–1596, doi:10.1038/s41588-021-00941-9 (2021).

11 Kaya-Okur, H. S., Janssens, D. H., Henikoff, J. G., Ahmad, K. & Henikoff, S. Efficient low-cost chromatin profiling with CUT&Tag. Nat Protoc 15, 3264–3283, doi:10.1038/s41596-020-0373-x (2020).

12 Peterson, C. L. & Workman, J. L. Promoter targeting and chromatin remodeling by the SWI/SNF complex. Curr Opin Genet Dev 10, 187–192, doi:10.1016/s0959-437x(00)00068-x (2000).

13 Kato, G. J., Barrett, J., Villa-Garcia, M. & Dang, C. V. An amino-terminal c-myc domain required for neoplastic transformation activates transcription. Mol Cell Biol 10, 5914–5920, doi:10.1128/mcb.10.11.5914-5920.1990 (1990).

14 Pellanda, P. et al. Integrated requirement of non-specific and sequence-specific DNA binding in Myc-driven transcription. EMBO J 40, e105464, doi:10.15252/embj.2020105464 (2021).

15 Krivtsov, A. V. & Armstrong, S. A. MLL translocations, histone modifications and leukaemia stem-cell development. Nat Rev Cancer 7, 823–833, doi:10.1038/nrc2253 (2007).

16 Crump, N. T. et al. MLL-AF4 cooperates with PAF1 and FACT to drive high-density enhancer interactions in leukemia. Nat Commun 14, 5208, doi:10.1038/s41467-023-40981-9 (2023).

17 Krivtsov, A. V. et al. A Menin-MLL Inhibitor Induces Specific Chromatin Changes and Eradicates Disease in Models of MLL-Rearranged Leukemia. Cancer Cell 36, 660–673 e611, doi:10.1016/j.ccell.2019.11.001 (2019).

18 Andrews, R. G., Singer, J. W. & Bernstein, I. D. Monoclonal antibody 12-8 recognizes a 115-kd molecule present on both unipotent and multipotent hematopoietic colony-forming cells and their precursors. Blood 67, 842–845 (1986).

19 Griffin, J. D., Ritz, J., Nadler, L. M. & Schlossman, S. F. Expression of myeloid differentiation antigens on normal and malignant myeloid cells. J Clin Invest 68, 932–941, doi:10.1172/jci110348 (1981).

20 Bennett, J. M. et al. Proposals for the classification of the acute leukaemias. French-American-British (FAB) co-operative group. Br J Haematol 33, 451–458, doi:10.1111/j.1365-2141.1976.tb03563.x (1976).

21 Khoury, J. D. et al. The 5th edition of the World Health Organization Classification of Haematolymphoid Tumours: Myeloid and Histiocytic/Dendritic Neoplasms. Leukemia 36, 1703–1719, doi:10.1038/s41375-022-01613-1 (2022).

22 Zentner, G. E., Tesar, P. J. & Scacheri, P. C. Epigenetic signatures distinguish multiple classes of enhancers with distinct cellular functions. Genome Res 21, 1273–1283, doi:10.1101/gr.122382.111 (2011).

23 Creyghton, M. P. et al. Histone H3K27ac separates active from poised enhancers and predicts developmental state. Proc Natl Acad Sci U S A 107, 21931–21936, doi:10.1073/pnas.1016071107 (2010).

24 Rada-Iglesias, A. et al. A unique chromatin signature uncovers early developmental enhancers in humans. Nature 470, 279–283, doi:10.1038/nature09692 (2011).

25 Chen, S., Zhou, Y., Chen, Y. & Gu, J. fastp: an ultra-fast all-in-one FASTQ preprocessor. Bioinformatics 34, i884–i890, doi:10.1093/bioinformatics/bty560 (2018).

26 Langmead, B. & Salzberg, S. L. Fast gapped-read alignment with Bowtie 2. Nat Methods 9, 357–359, doi:10.1038/nmeth.1923 (2012).

27 Li, H. et al. The Sequence Alignment/Map format and SAMtools. Bioinformatics 25, 2078–2079, doi:10.1093/bioinformatics/btp352 (2009).

28 Quinlan, A. R. & Hall, I. M. BEDTools: a flexible suite of utilities for comparing genomic features. Bioinformatics 26, 841–842, doi:10.1093/bioinformatics/btq033 (2010).

29 Kent, W. J., Zweig, A. S., Barber, G., Hinrichs, A. S. & Karolchik, D. BigWig and BigBed: enabling browsing of large distributed datasets. Bioinformatics 26, 2204–2207, doi:10.1093/bioinformatics/btq351 (2010).

30 Robinson, J. T. et al. Integrative genomics viewer. Nat Biotechnol 29, 24–26, doi:10.1038/nbt.1754 (2011).

31 Ramirez, F. et al. deepTools2: a next generation web server for deep-sequencing data analysis. Nucleic Acids Res 44, W160–165, doi:10.1093/nar/gkw257 (2016).

32 Meers, M. P., Tenenbaum, D. & Henikoff, S. Peak calling by Sparse Enrichment Analysis for CUT&RUN chromatin profiling. Epigenetics Chromatin 12, 42, doi:10.1186/s13072-019-0287-4 (2019).

33 Li, Q. H., Brown, J. B., Huang, H. Y. & Bickel, P. J. Measuring Reproducibility of High-Throughput Experiments. Ann Appl Stat 5, 1752–1779, doi:10.1214/11-Aoas466 (2011).

34 Neph, S. et al. BEDOPS: high-performance genomic feature operations. Bioinformatics 28, 1919–1920, doi:10.1093/bioinformatics/bts277 (2012).

35 Machanick, P. & Bailey, T. L. MEME-ChIP: motif analysis of large DNA datasets. Bioinformatics 27, 1696–1697, doi:10.1093/bioinformatics/btr189 (2011).

36 Grant, C. E., Bailey, T. L. & Noble, W. S. FIMO: scanning for occurrences of a given motif. Bioinformatics 27, 1017–1018, doi:10.1093/bioinformatics/btr064 (2011).

